# NMR structure of the Orf63 pro-lytic protein from lambda bacteriophage

**DOI:** 10.1101/2023.10.03.560691

**Authors:** Naushaba Khan, Tavawn Graham, Katarzyna Franciszkiewicz, Sylwia Bloch, Bożena Nejman-Faleńczyk, Alicja Wegrzyn, Logan W Donaldson

## Abstract

The *orf63* gene resides in a region of the lambda bacteriophage genome between the *exo* and *xis* genes and is among the earliest genes transcribed during infection. In lambda phage and Shiga toxin (Stx) producing phages found in enterohemorrhagic *E. coli* (EHEC) associated with food poisoning, Orf63 expression reduces the host survival and hastens the period between infection and lysis thereby giving it pro-lysogenic qualities. The NMR structure of dimeric Orf63 reveals a fold consisting of two helices and one strand that all make extensive intermolecular contacts. Structure-based data mining failed to identify any Orf63 homolog beyond the family of temperate bacteriophages. A machine learning approach was used to design an amphipathic helical ligand that bound a hydrophobic cleft on Orf63. This approach may open a new path towards designing therapeutics that antagonize the contributions of Stx phages in EHEC outbreaks.

## Introduction

*E. coli* O157:H7 has been associated with food- and water-borne infections throughout the world. While only a few percent of affected people are hospitalized, the impact can be large when infection leads to a community outbreak. Since the discovery of *E. coli* O157:H7 over forty years ago^1^, other clinically important strains have emerged^2^. One way in which pathogenic *E. coli* are genetically distinct from their benign counterparts is that their genome contains prophage DNA which bears *stx* genes, encoding Shiga toxins. Such a phage is called Shiga toxin-converting phage, or Stx phage, and as a prophage it remains dormant until a stress pathway is activated in the host bacteria leading to a developmental transition from a lysogenic state to an activated lytic state. As the phage replicates, it also produces one of two forms of the Shiga toxin enzyme (Stx1 or Stx2,) that enters intestinal epithelial cells and kills them due to ribosome inactivation^3^. While the oxidative attack by intestinal immune cells offers one way to induce a stress response in bacteria, antibiotic treatment may also lead to the same outcome. As a result, antibiotics that would normally be the first treatment against a bacterial infection are avoided during a Shiga-toxin producing *E. coli* (STEC) infection. This critical limitation impresses a need to explore phage-bacteria dynamics further and identify new therapeutic stratagies^4,5^.

Stx phages share many characteristics with phage λ, a model system associated with many landmark discoveries in molecular biology. One shared genetic region found between the *exo* and *xis* genes encodes five proteins (Orf60a, Orf61, Orf63, Orf73, and Ea22) that may work separately or together at the earliest stages of infection to affect the commitment to a lysogenic or lytic developmental outcome^6–8^ (Fig. 1). Originally, *exo-xis* genes were associated with cell cycle and DNA replication changes driven by the *p*_L_ promoter^9^ during the earliest stages of infection with phage λ^10^. Assays that measure the period between induction and the observation of new viral progeny, the number of bacterial survivors after infection, and the efficiently in which bacteria are converted to lysogens all suggest that Orf60a, Orf61, and Orf63 are pro-lytic where Ea22 and Orf73 tend to be pro-lysogenic^8,11^. Orf63, the subject of this report, is a 63 amino acid protein that is highly conserved among λ and Stx phages alike^12^ Since the effects of Orf63 and other *exo-xis* region gene products were amplified in STEC bacteria relative to benign *E. coli* strains infected with λ^13^, a closer examination may reveal new protein partnerships and pathways that Stx phages use to favor their success within their host.

**Figure 1.**
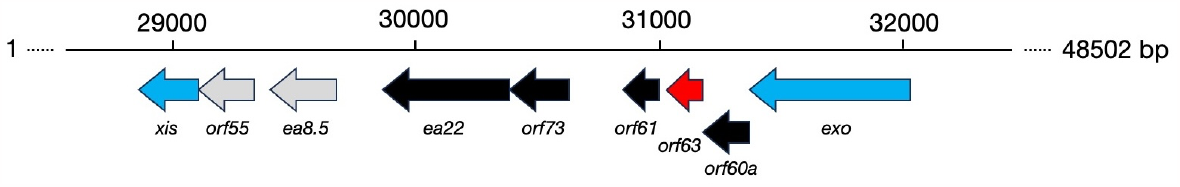
The *exo-xis* region of phage λ. This region may be considered in terms of a conserved and largely invariant region consisting of *orf60a, orf63* (red), *orf61, orf73*, and *ea22* followed by a region with a variable gene composition (grey).

Here, we present the structure of Orf63 as a first step towards understanding its functional role. This investigation increases the repertoire of *exo-xis* proteins that have either been modeled or solved to date^14,15^. Through molecular modeling, we have identified a region on Orf63 that can bind a short α-helical ligand. Protein partners of Orf63 may share a similar binding mode. Orf63 dimerizes in a way that has not been previously observed in the Protein Data Bank. All predicted structures by machine learning methods in comprehensive protein databases all point to proteins that share Orf63 signature sequences, including the preservation of an intermolecular ionic bond.

## Results

Orf63 is a small protein that is known as DUF1382 in the Conserved Domain Database (CDD) and as IPR009814 in the InterPro protein family database as IPR009814. Size exclusion chromatography multi-angle laser scattering (SEC-MALS) revealed that Orf63 occurs exclusively as a dimer in solution (Supplementary Fig. S1). The Orf63 protein was the most soluble and produced the highest quality NMR spectra at pH 7.5 with little dependence on ionic strength. Following this preliminary survey, the NMR structure of 6xHis-Flag affinity tagged Orf63 was determined using a conventional combination of experiments supplemented with a ^12^C-^13^C filtered-edited NOESY experiment to obtain intermolecular contacts. We refer the reader to Supplementary Fig. S2 for selected data from ^12^C-^13^C filtered edited NOESY and ^13^C-filtered NOESY spectra corroborating the hydrophobic core and secondary structures. A total of 796 distance restraints, 25 hydrogen bond restraint pairs, and 30 torsion angle restraints were used as input for a two-stage structure calculation that used CYANA to produce an initial ensemble and Rosetta for final refinement. A complete statistical summary of the structure determination is provided in Supplementary Table S1. Overall, the RMSD for ordered regions of the ensemble of the 20 lowest energy structures was 0.8 Å for backbone atoms and 1.1 Å for all heavy atoms. The backbone RMSD between the NMR structure and the AlphaFold model was 1.8 Å.

The structural features of λ Orf63 are summarized in Fig. 2. From the initial assignments, 12 amino acids from the N-terminus and 11 amino acids from the C-terminus were determined to be unstructured leaving a very compact dimeric fold consisting of two α-helices and one β-strand in the remaining 40 amino acids. The tightly intertwined dimer can be folded by placing two extended chains beside each other to form the anti-parallel, two-stranded β-sheet is formed first, and then folding the two α-helices above and below the β-sheet. The dimer interface draws contributions from much of the protein making it unlikely that the protein would ever be observed in a monomeric state. A survey of the PDB using SSM^16^ and FoldSeek^17^ revealed no homologous fold. Extending the survey to large databases who structures have been predicted by AlphaFold, the only homologous examples were other viral Orf63 proteins. The β-strand extends includes the sequence 28-FVPIP-32. The two prolines are positioned in the β-sheet among a regular series of hydrogen bonds supported by backbone HN-HN, HN-HA, and HA-HN NOE observations. In summary, the structure of Orf63 presents a newly identified fold and mode of dimerization that is exclusive to the λ family of bacteriophages.

**Figure 2.**
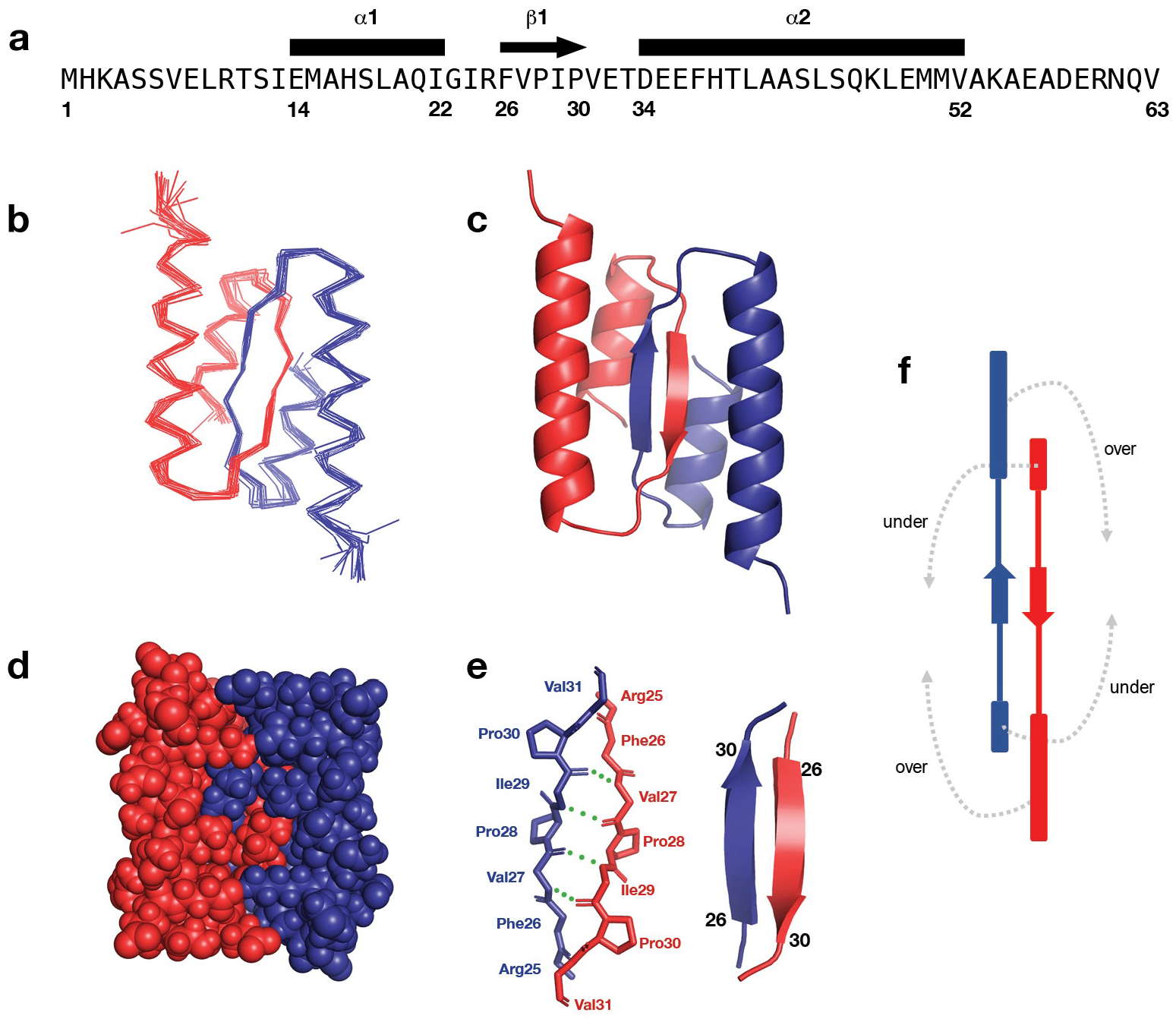
Structural characteristics of λ Orf63. **(a)** Observed secondary structural elements mapped onto the sequence. **(b)** A Cα trace of the best 20 structures in the ensemble **(c, d)** A cartoon and space filling representation of the dimer **(e)** Hydrogen bond network across the β-strands at the dimer interface. **(f)** Topology diagram of the dimer. The fold may be considered in terms of the two a-helices folding under and over the β-sheet.

Variation among a set of 290 Orf63 homologs in the UniRef100 database is presented in Fig. 3. As expected, most of the conserved amino acids mapped to the hydrophobic core of the protein. The central β-strand bridge was intolerant to variation at positions corresponding to F28, V29, and P32 in λ Orf63. Two arginines (R3/R10) that found in the unstructured N-terminal region appear to be conserved with unknown significance. Two ionic bonds observed in the structure are absolutely conserved (E36/R47) in one case and partially conserved (R25/D34) in another case. With respect to the latter case, the chemical shifts corresponding to the backbone of D34 could not be assigned suggesting that it was dynamic in the intermediate timescale at the beginning of the second helix. As a result, the corresponding ionic bond identified in the structure is likely to be poorly defined, as well.

**Figure 3.**
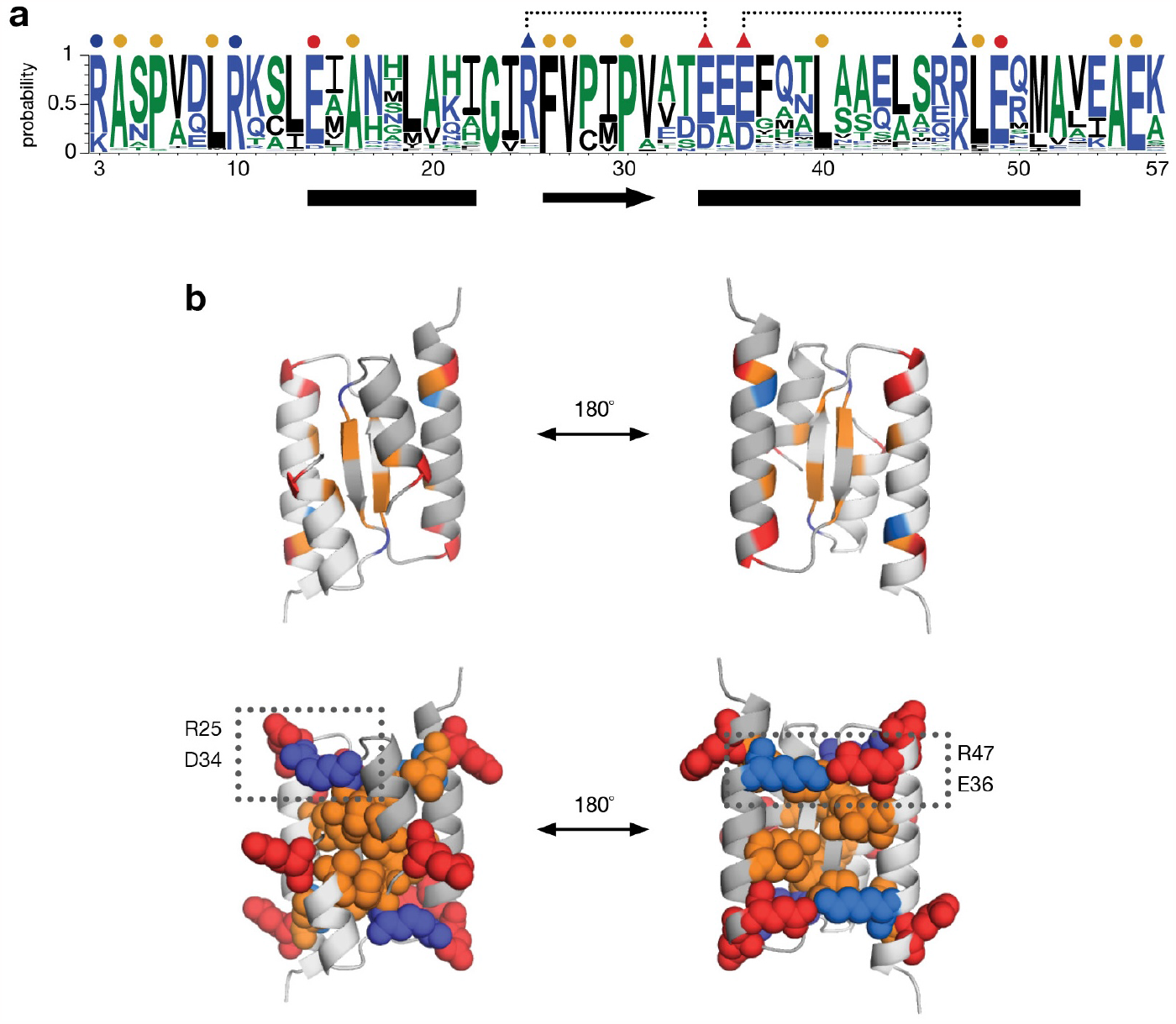
Sequence variation among Orf63 protein in the UniRef100 database. **(a)** Above the Weblogo consensus plot, a dot at nonpolar (orange), acidic (red), and basic positions (blue) indicates conservation >80%. Triangles indicate positions where an ionic bond was observed in the structure. **(b)** Cartoon and space filling views of the most conserved residues with the indicated ionic bonds.

We hypothesized that if the E36K/R47 ionic bond was an essential feature of the fold, breaking the ionic bond would have profound effects on the structure and stability of protein. To answer this question, a variant Orf63[E36K] was made and was assessed by circular dichroism (CD) spectroscopy along with a sample of wild type protein. As shown in Fig. 4, the far-UV CD spectrum Orf63[E36K] suggested a potential loss of structure corroborated by the absence of two strong α-helical signals at 210 nm and 222 nm. While a thermal denaturation assay monitored at multiple wavelengths revealed a Tm of ∼75 °C for wild type Orf63, no transition was observed for Orf63[E36K] suggesting it was unfolded at all temperatures surveyed. Wild type Orf63 chemical denatured at a ∼2.5 M concentration of urea. Both the thermal and chemical denaturation of wild type Orf63 was reversible.

**Figure 4.**
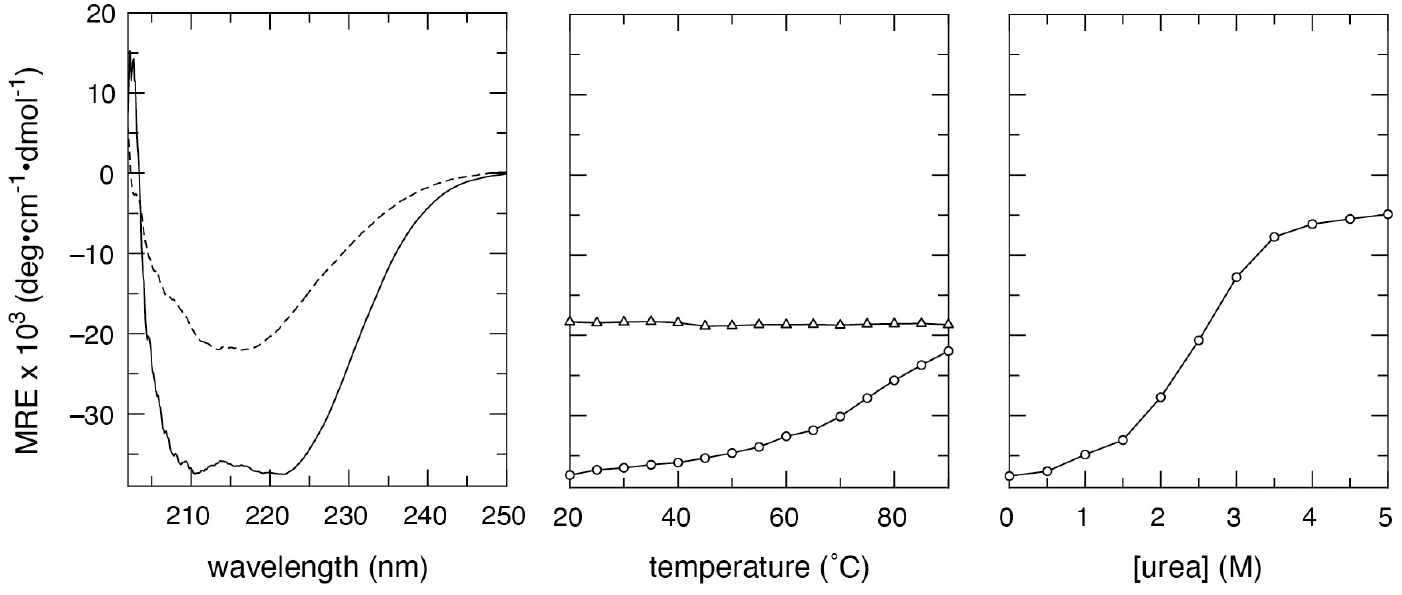
CD spectroscopy. (a) Far UV spectrum of wild type Orf63 (solid line) and Orf63[E36K] (dashed line). (b) Thermal denaturation of Orf63 (circles) and Orf63[E36K] (triangles) monitored at 222 nm. (c) Urea denaturation of wild type Orf63 (circles).

*E. coli* endogenously producing either wild type Orf63 or an Orf63 variant demonstrated different degrees of survival. Comparing the white bars in Fig. 5, wild type Orf63 reduced bacterial by nearly an order of magnitude compared to the unfolded and presumably non-functional E36K variant. Other variants explored solvent-facing positions on helix α2. The Q46E and K54E variants appear to abolish Orf63 function whereas the H38E variant retained the functional characteristics of the WT protein. Orf63 and the tested variants that negatively affect bacterial survival also promoted a stronger lytic outcome after infection with phage λ mutant lacking *orf63* (black bars in Fig. 5). Since the number of lysogens observed after infection was comparable to the number of bacterial survivors reinforcing the role of Orf63 appears to operate exclusively in lytic developmental pathways.

**Figure 5.**
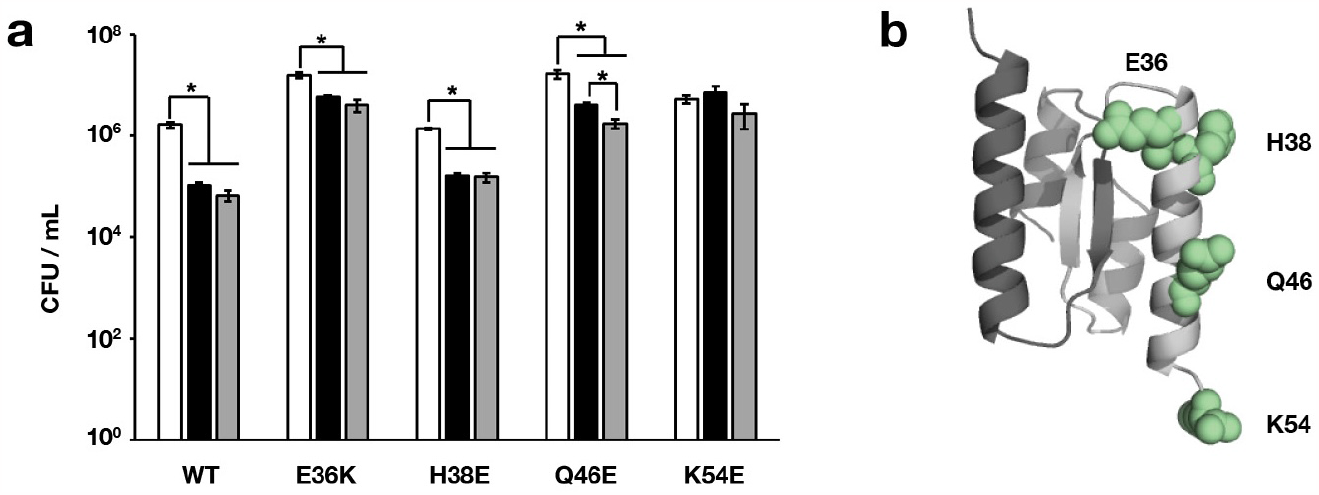
Effect of Orf63 and four variants on host survival and viral development. **(a)** Phage λΔ*orf63* at an m.o.i. of 5 was added to *E. coli* C600 previously induced with IPTG to express different variants of the *orf63* gene and to produce different forms of the Orf63 protein. White bars represent bacteria that were mock infected with buffer as a negative control. Black bars represent bacterial survivors post-infection. Grey bars represent lysogens among bacterial survivors. All counts are expressed as the number of colony forming units (CFU) per mL. Errors are presented as SD from three replicates. **(b)** Location of the variants examined on the structure of Orf63.

A bioinformatics survey of performed on a set of domains of unknown functions (DUFs) suggested that Orf63 may have nucleic acid binding properties^18^. Our observation that polar, solvent facing Orf63 variants on helix α2 affected activity, a sample of ^15^N-Orf63 with a molar excess of a 32 bp palindromic dsDNA that was originally determined to be a high-affinity binding partner for the human Sox9 DNA binding domain^19^. A comparison of ^1^H-^15^N HSQC spectra of free and DNA-bound Orf63 showed no chemical shift changes or line broadening suggesting that in this limited context, Orf63 may require a specific DNA sequence to achieve high affinity binding or not bind dsDNA, at all.

A yeast two-hybrid study of phage λ proteins^20^ identified the transcription factor YqhC as a protein partner of Orf63. YqhC regulates the production of the NADPH-dependent aldehyde reductase YqhD known for its ability to confer resistance to glutaraldehyde, a commonly used disinfecting agent^21^. Highly purified YqhC was prepared and desalted into the same buffer as a preparation of ^15^N-Orf63. ^1^H-^15^N HSQC spectra were acquired for 2:1 mixture of YqhC:^15^N-Orf63 and a ^15^N-Orf63 control. Upon comparison, no chemical shift changes were observed suggesting that YqhC is not a protein partner of Orf63 (Supplementary Fig. S3).

If Orf63 has the potential to bind another protein, a hydrophobic cleft formed between helix α1 and α2 emerges an interesting candidate to explore (Fig. 6). Using a machine learning approach based on AlphaFold, a single helix was docked and sampled to identify the sequence that presents the most complementary contacts to Orf63. The candidate from this procedure was further sampled with Rosetta to reveal the most important sites on the helical ligand and how much they could be varied. The best helical candidate was selected on the basis of most favorable energy. Since the N-terminus of the helical ligand was close to the C-terminus of Orf63, a hybrid protein was designed with a short intervening linker. This hybrid protein was then used as input to AlphaFold to verify that it predicted the same fold as the complex. The ^1^H-^15^N HSQC spectrum of the hybrid protein resembled Orf63 with additional peaks from the helical ligand along with a set of chemical shift changes likely arising from structural differences at the Orf63-ligand interface. To confirm the specificity of the interaction, the same 20-mer sequence was expressed as a 6xHis-tagged-Ubquitin fusion protein. A 2:1 mixture of the Ubiquitin fusion protein to with ^15^N-labelled Orf63 dimer revealed significant line broadening suggesting binding was occuring (total molecular weight of the complex was 45 kDa). In contrast, a 2:1 mixture of a Ubiquitin fusion protein with an unrelated sequence produced no chemical shift or line broadening changes (Supplementary Fig. S4). While the NMR data provides evidence that the designed peptide is ligand of Orf63, our attempt at determining affinity by biolayer interferometry failed suggesting the the K_d_ of the designed peptide peptide for Orf63 is poor (>20 μM).

**Figure 6.**
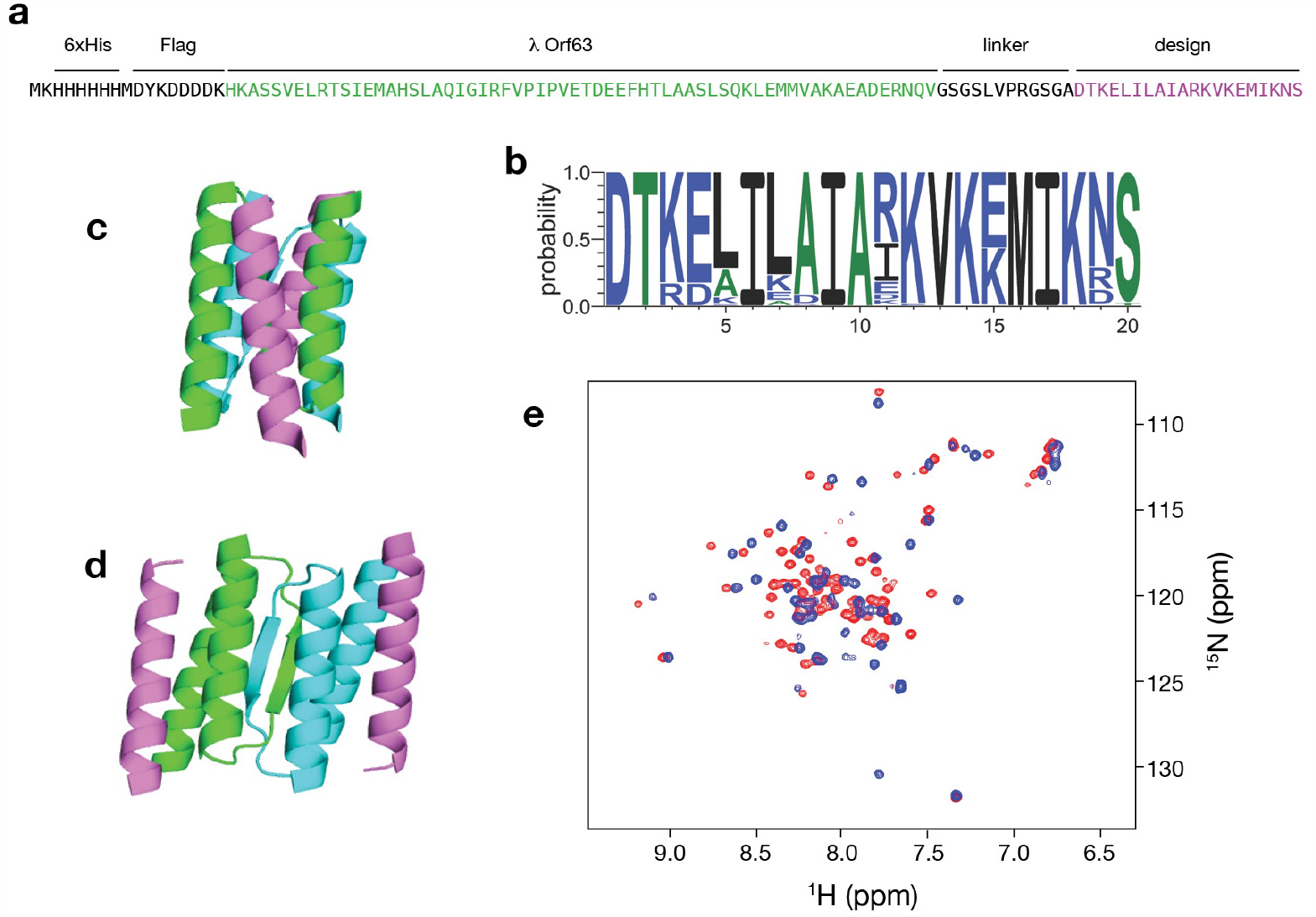
A designed α-helix binds a cleft made by helix α1 and α2 of Orf63. **(a)** A hybrid Orf63 with the designed α-helix appended to the C-terminus. **(b)** Sequence variation throughout the designed helix sampled by Rosetta **(c, d)** Cartoon representations of the designed helix (purple) against each Orf63 protomer (green/blue). **(e)** A ^1^H-^15^N HSQC spectrum of a hybrid Orf63 protein (red) reveals a set of new peaks and chemical shift differences relative to the spectrum of Orf63 (blue).

## Discussion

Orf63 is a non-essential dimeric protein in both λ and related Shigatoxigenic phages that promotes lytic development over lysogenic development. The *orf63* gene is located within a cluster of conserved, yet under-characterized open reading frames found in the *exo-xis* region of the phage genome. From sequence and structural datamining, Orf63 is unique to temperate phages. Despite its small size and potential to act as a modular domain, it is autonomous and not part of any protein.

The NMR structure of Orf63 reveals a unique fold with a dimer interface drawing contributions from each of the two α-helices and one β-strand. An essential ionic bond between E36 and R47 reinforces the dimer interface. Unexpectedly, there is sequence conservation of several charged amino acids in N-terminal and C-terminal unstructured segments flanking the ∼40 amino acid domain. The role of these flanking amino acids is unknown but it is tempting to speculate that they may be involved in protein networks by donating small linear motifs^22^ to a protein partner or making contacts with nucleic acids. A mass spectrometry-based proteomics survey is an obvious experimental route toward identifying Orf63 partners either from viral or bacterial origins.

From bacterial survival and lysogenic conversion assays, solvent exposed amino acids in helix α2 appear to modulate activity. The nature of the potential contacts they make may be augmented by a deep hydrophobic cleft formed by helix α1 and α2. A more exhaustive mutagenic survey is required to map a functional surface patch on Orf63. Taking a different approach towards this objective, a helical ligand was designed that could bind the α1/α2 cleft as shown by differences in NMR spectra between Orf63 and a hybrid Orf63 with the ligand fused to the C-terminus. One helix alone as a ligand creates a potential for a variety of host and viral proteins to partner with Orf63. After this study was performed, the machine learning software RFdiffusion was released enabling larger binding proteins to be designed^23^. Despite requesting a binding protein of 80-100 amino acids to the software, the same short helical ligand was repeatedly observed in the output thereby reinforcing our initial observation that one helix would be sufficient. The ability to design antagonistic binding peptides and proteins offers new way to create new therapeutics against EHEC outbreaks.

## Methods

### Expression and purification of proteins

Phage λ *orf63* (laboratory reference #201916, UniProt: Q38267; DUF1382) was gene synthesized by ATUM (Menlo Park, CA) with an amino-terminal 6xHis + FLAG tag and inserted into plasmid pD441NH (T5 promoter). Amino acid substitution mutants were prepared using the Q5 mutagenesis method (New England Biolabs) with oligonucleotides from IDT. Sequencing was performed by The Centre for Applied Genomics (Hospital for Sick Children; Toronto, ON) Expression was achieved in cultures under 50 μg/mL kanamycin selection at 30 °C in an *E. coli* BL21. At an A600 of 0.8, expression was induced by the addition of 1 mM isopropyl thiogalactoside (IPTG), and the culture was grown for 3 h further before harvesting. Detailed information regarding the purification of this fragment by nickel affinity and gel filtration chromatography has been published previously^13^. For NMR spectroscopy, a uniformly ^13^C, ^15^N isotopically labelled protein sample was made from a 1 L culture containing M9 minimal media salts supplemented with 3 g of ^13^C-glucose (Sigma-Aldrich), 1 g of ^15^N ammonium chloride (Sigma-Aldrich) and 1 g of ^13^C,^15^N Celtone algal extract (Cambridge Isotopes). Natural abundance Orf63 proteins were prepared using a similar protocol except cultures were grown in LB media. A potential protein partner of Orf63, YqhC, identified from a previous yeast two-hybrid study was gene synthesized by Gene Universal (Newark, NJ) and inserted into the *NdeI*/*XhoI* multiple cloning sites of pET29b (Novagen) leading to a carboxy-terminal 6xHis tagged protein. A 1 L culture of *E. coli* BL21:DE3 in LB media under 50 μg/mL kanamycin selection was incubated at 37 °C until an A600 of 0.5 was reached, then the culture was chilled, protein expression was induced with 1 mM ITPG, and the culture was incubated further for 18 h at 16 °C. Protein purification followed a similar two-step nickel-affinity / gel-filtration protocol used for Orf63. Protein concentrations were estimated by UV absorbance at 280 nm. Two ubiquitin fusion proteins with an N-terminal 6xHis tag, linker GGLVPRGSG, and either the sequence DTKELILAIARKVKEMIKNS corresponding to the designed Orf63 binding peptide (pET28-UbqOrf63L2; reference #7A03005 or an unrelated peptide MAIAHAATEYVFADFVLK (pET28-UbqPanxL1; reference #7A03212-2) were expressed in *E. coli* BL21:DE3 at 37 °C, and purified according to the two step nickel-affinity / gel-filtration protocol. The final buffer for all expressed protein was either NMR bufffer (5 mM Tris-Cl pH 7.4, 50 mM NaCl, 0.05% NaN_3_) or phosphate buffered saline (PBS) pH 7.4.

### NMR sample preparation

A preparation of λ Orf63 protein was concentrated to 0.8 mM in NMR buffer supplemented with 10% (v/v) D_2_O for NMR spectroscopy. For NMR experiments requiring a sample in D_2_O, the Orf63 protein was dialyzed to a similar buffer in 98% (*v/v*) D_2_O. A mixed ^12^C/^13^C sample was made by mixing ^12^C/^14^N and ^13^C/^15^N proteins at a 1:1 molar ratio, adding urea to 6 M and rapidly diluting the mixture into a 20-fold excess of NMR buffer. The protein was concentrated and dialyzed to a 98% (*v/v*) D_2_O buffer.

### NMR spectroscopy

A series of 2D and 3D heteronuclear NMR spectra were acquired at a temperature of 308 K on a 700 MHz Bruker AvanceIII spectrometer equipped with a nitrogen-chilled probe to support the backbone assignments that had been previously described. These experiments included (2D-15N-HSQC, 2D-13C-HSQC, 3D-CCONH, 3D-HCCONH, 3D-HBHACONH, and 3D-CCH-TOCSY). The 3D experiments were acquired according to a 10-20% sparse sampling schedule and processed with NMRPipe^24^ and HMSist^25^. Distance restraints were obtained from a 3D-15N-NOESY and a 3D-13C-NOESY experiments sparsely sampled at 20%. Intermolecular distance restraints were obtained from a 3D-^12^C-filtered/^13^C-edited NOESY experiment acquired at the laboratory of Lewis Kay (Univ. Toronto) on a Varian Inova 600 MHz instrument equipped with a room temperature probe.

### Structure determination

NMR spectra were analyzed with CCPN Analysis 2.52^26^. Backbone dihedral angles were predicted from chemical shifts with TALOS^27^. An initial ensemble of 100 structures sorted by the lowest number of NOE violations were calculated from a set of 10000 structures using CYANA 3^28^. The ensemble was refined using Rosetta with distance, angle and hydrogen bond restraints converted to Rosetta .cst format. Symmetry was enforced throughout the calculation. Details of the Rosetta refinement method have been published^29^. The top 20 refined structure solutions that satisfied the experimental restraints the most formed the final ensemble. Structural quality was assessed with PSVS^30^ and PROCHECK^31^.

### Determination of oligomeric state

Size-exclusion chromatography multi-angle laser scattering (SEC-MALS) was performed at the Hospital for Sick Children SPARC analytical facility (Toronto, ON) on a 1260 Infinity-II HPLC (Agilent) and SEC300A 2.7 μM 4.6×300mm column (Advance Bio) analytical column at 0.2 mL/min linked to MiniDawn TREOS MALS and a OptiLab T-rEX refractive index (RI) detection instruments (Wyatt). The chromatography and detection system were equilibrated in 10 mM Tris-Cl, 0.15 M NaCl, pH 8.0 for 18 h before the first detection. A preliminary injection of 2 mg/mL of bovine serum albumin was performed to calibrate peak and retention time profiles of the detectors prior to sample analysis. For the primary Orf63 analysis, a 2 mg/mL sample was centrifuged at 13,000 g at 4 °C for 10 min to remove any precipitates and other large, insoluble particles and applied to the system. Chromatogram data was analyzed with ASTRA (Wyatt) with a *dn/dc* of 0.185.

### Bioinformatics and structure predictions

The UniRef100 database^32^ was searched with mmseqs2^33^ for homologs to λ Orf63. Any truncated sequences were removed. The dataset was realigned in AliView^34^ and exported as FASTA formatted list to WebLogo3^35^. AlphaFold Multimer was used to predict the dimeric structure of Orf63 (*https://github.com/sokrypton/ColabFold*).

### Binding protein design

AfDesign was used to predict five 20-residue binding peptides for Orf63 (*https://colab.research.google.com/githum/sokrypton/ColabDesign/blob/main/af/examples/peptide_binder_design.ipynb)*. To assist modeling, a binding site on Orf63 was declared as a set of hotpot residues. The best design docked to Orf63 was used as input to Rosetta *fixbb* design with all amino acid positions being allowed to vary. From an ensemble of 512 structure a consensus sequence was determined. The sequence with the highest number of observations was selected. A hybrid Orf63 protein was designed by appending the sequence of best design to the C-terminus of Orf63 separated by the linker, GSGSLVPRGSGA. A codon optimized gene corresponding to this design was synthesized by Gene Universal (Newark, DE) and inserted into pET29b. Expression and purification were achieved by growing a culture in M9 media for ^15^N-labeling at 37 °C and following the previously described protocol for induction of gene expression and protein purification.

### Bacterial survival assay

To stimulate the production of Orf63 protein or its derivatives from the pD441NH plasmid, overnight bacterial cultures of *E. coli* C600 were diluted 100-fold in a fresh LB medium and treated with 1mM IPTG. Host bacteria were cultivated at 37°C to A600 of 0.1. A 0.1 mL aliquot was centrifuged (2,000 *g* for 10 min at 4°C). and afterwards, the pellet was washed with TCM buffer (10 mM Tris-HCl, 10 mM MgSO4, 10 mM CaCl_2_, pH 7.2) twice, and then resuspended in the same buffer. Bacteriophage λΔ*orf63* was added to bacteria at an m.o.i. of 5. The mixture was incubated at 37°C for 15 min and afterwards, serial dilutions in 0.85% NaCl were prepared. From these dilutions, 40 μL of each suspension was spread on LB agar plates supplemented with 25 μg/mL kanamycin. After an overnight incubation at 37°C, the number of *E. coli* survivors per mL (CFU/ml) was counted.

### Lysogenic conversion of survivors

Fresh bacterial colonies from an appropriate dilution were placed in a 96-well plate filled with 200 μL of LB medium and shaken at 37°C until the cultures reached an A600 of 0.1. The cultures were then treated with 50 J/m^2^ UV light to induce phage production and incubated at 37°C for 2 h further. The lysogens in the sample were mixed with chloroform and centrifuged a 2,000 *g* for 10 min at 4°C. The aqueous phase was spotted onto double-layer LB agar. After overnight incubation at 37°C, the number of lysogens on each agar plate was determined.

## Acknowledgements

This work was supported by National Science Center (Poland) within a project grant (UMO-2013/09/B/NZ2/02366) to A.W., and a Canadian NSERC Discovery Grant (2018-05838) to L.W.D.

## Author contributions statement

L.W.D. performed the sequence analyses and molecular modeling, acquired the NMR and CD data, and solved the NMR structure. N.K. and T.G. made proteins and mutants. K.F., S.B. and B.N-F. performed the phage development assays. LWD wrote the manuscript and B.N.-F. and A.W. revised the manuscript.

## Additional information

### Data deposition

Chemical shifts of λ Orf63 were deposited in the BMRB (entry 59519). The final ensemble of structures was deposited in the PDB (entry 8DSB).

### Supplementary information

A separate datafile includes Table S1 and Figures S1-S4.

