## Supplementary Information for "NMR structure of the Orf63 pro-lytic protein from lambda bacteriophage"

**Table S1.** Structural statistics for the Orf63 NMR structure

| | |
|---|---|
| NOE distance restraints in the ensemble[a] | 796 |
| intraresidue | 373 |
| short (\|*i*-*j*\| = 1) | 163 |
| medium (1 ≤ \|*i*-*j*\| ≤ 5) | 73 |
| long (\|*i*-*j*\| > 5) | 113 |
| interchain | 74 |
| Hydrogen bond distance restraints | |
| HN–O / N–O pairs | 25 |
| Torsion angle restraints | |
| backbone (Φ / Ψ) | 30 |
| Structural quality analysis | |
| close contacts | 0 |
| RMS deviation of bond angles (deg) | 0.3 |
| RMS deviation of bond lengths (Å) | 0.0009 |
| RMS deviation to the mean coordinates[b] | |
| all backbone / heavy atoms (Å) | 0.8 / 1.1 |
| Ordered backbone / heavy atoms (Å) | 0.5 / 0.8 |
| Ramachandran plot[c] (%) | |
| residues in most favored regions | 98.8 |
| residues in additional allowed regions | 1.2 |
| residues in generously allowed regions | 0.0 |
| residues in disallowed regions | 0.0 |

[a] None of the twenty structures in the ensemble (PDB: 8DSB) has a distance violation > 0.2 Å and a dihedral angle violation > 5°.

[b] Ordered residues (16-54) are defined by a dihedral angle order parameter with S(Φ)+S(Ψ)>=1.8 as determined by PSVS.

[c] Determined by PROCHECK.

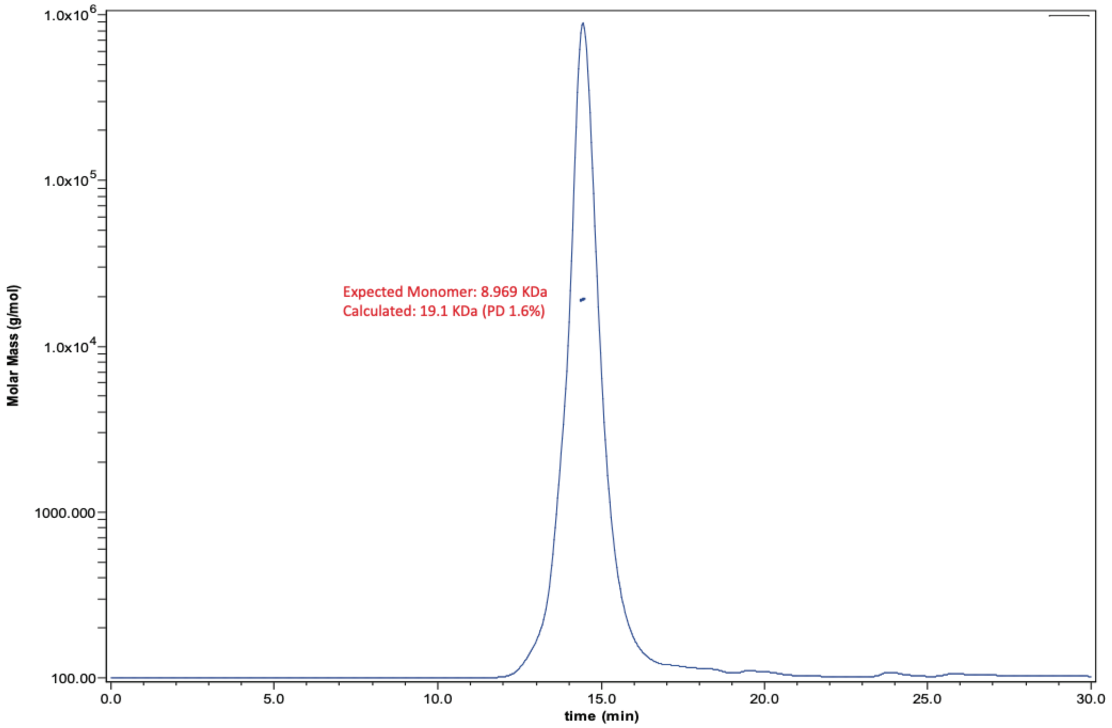


**Figure S1** — Size exclusion chromatography - multiangle laser scattering analysis (SEC-MALS) of a 2 mg/mL injection of λ Orf63. Analysis of the primary peak demonstrates that the protein is dimeric in solution.

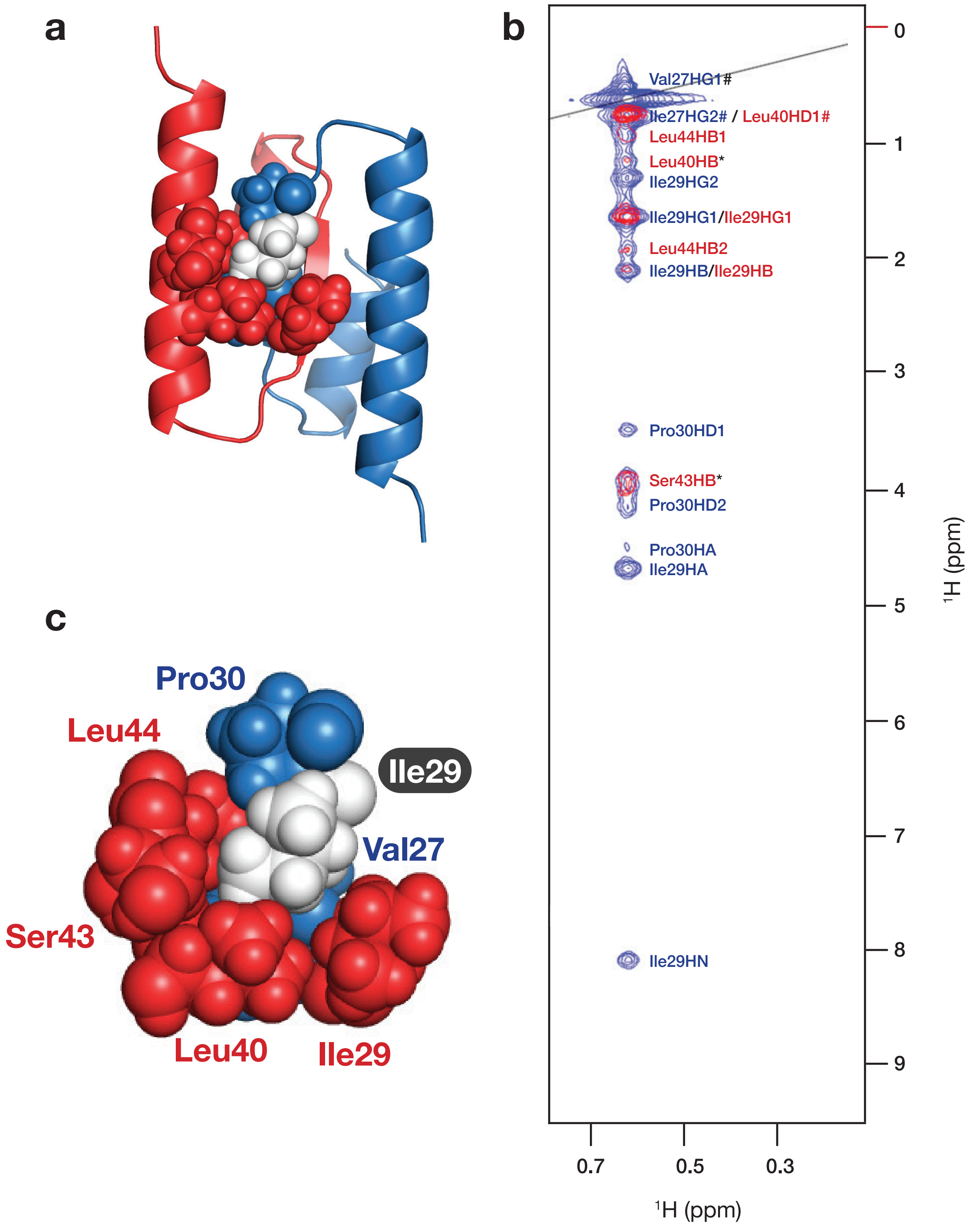


**Figure S2** — NOE observations supporting the structure of Orf63. **(a,b)** Cartoon diagram and magnification of intramolecular amino acids (blue) and intermolecular amino acids (red) in the vicinity of Ile43 (white). **(c)** A representative plane from a 3D $^{13}$C-edited NOESY spectrum (blue) and 3D $^{12}$C-filtered,$^{13}$C-edited NOESY spectrum (red) highlighting NOEs from the HD1 methyl group of Ile29.

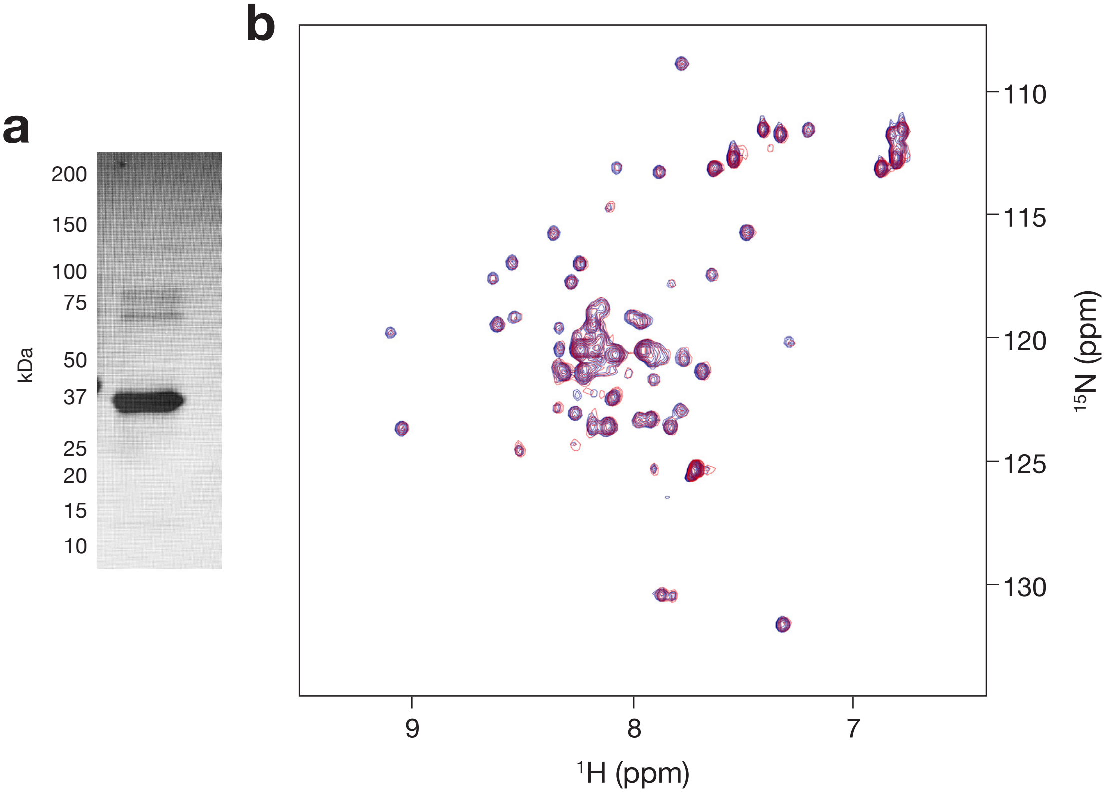


**Figure S3** — λ Orf63 does not bind YqhC. **(a)** A sample of 6xHis-tagged *E. coli* YqhC was purified by nickel affinity and gel filtration chromatography. The sample was assesed to be very pure by SDS-PAGE. YqhC protein was mixed at a 2:1 ratio with $^{15}$N-Orf63 in the same buffer (phosphate buffed saline, pH 7.4, 10% $D_2O$) and concentrated to 0.15 mM. **(b)** An $^{1}$H-$^{15}$N HSQC spectrum of $^{15}$N-Orf63 (blue) and $^{15}$N-Orf63:YqhC (red).

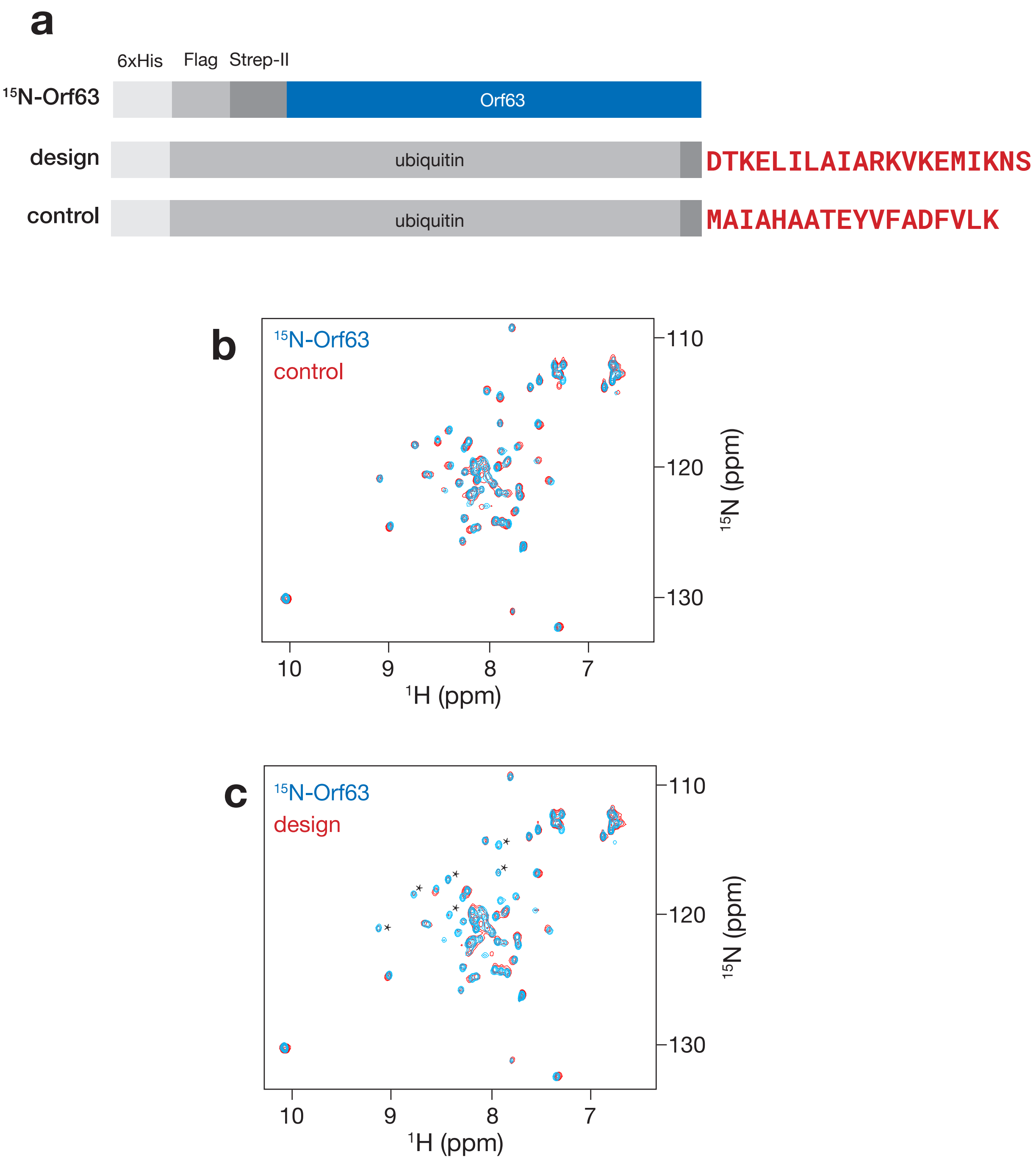


**Figure S4** — A designed peptide binds Orf63. **(a)** A mixing experiment was performed with $^{15}$N-Orf63 and a Ubiquitin-fusion protein with a C-terminal extension corresponding to a designed peptide, or an unrelated peptide control. The mixing ratio was 2:1 (Ubiquitin fusion to Orf63 dimer) in 5 mM Tris-Cl pH 7.4, 50 mM NaCl. **(b,c)** $^{15}$N-HSQC spectra of $^{15}$N-Orf63 (blue) and $^{15}$N-Orf63 mixed with a Ubiquitin fusion protein. Asterisks denote peaks that were severely line broadened in the mixing experiment.
